# Two decades of compositional restructuring of soil biodiversity in Germany despite stable α- and β-diversity indices

**DOI:** 10.1101/2025.07.21.665925

**Authors:** Judith Paetsch, Juliane Romahn, Kathrin Theissinger, Damian Baranski, Lena Bonassin, Leonie Schardt, Jan Koschorreck, Henrik Krehenwinkel, Miklós Bálint

## Abstract

Soil ecosystems host some of the most taxonomically and functionally diverse biological communities on Earth, yet long-term trends in their biodiversity remain poorly understood. Here, we analysed soil biodiversity dynamics over 20 years with samples archived in the German Environmental Specimen Bank. We assessed temporal and spatial patterns in α-diversity and β-diversity with shotgun metagenomics across bacteria, fungi, and metazoa. We found no statistically significant temporal trends in α-diversity for any group. Total β-diversity also appeared temporally stable. However, decomposing β-diversity into its balanced variation and abundance gradients revealed taxon-specific compositional restructuring. Bacterial and fungal communities showed signs of compositional homogenisation, while metazoan communities remained more stable. Spatial structuring was pronounced across all groups. Land use emerged as a key spatial predictor of community composition for bacteria and fungi, and geographic locality for metazoans. Our findings show that apparent stability in standard biodiversity indices may mask significant underlying community change. This highlights the need for integrative, taxonomically inclusive approaches to biodiversity monitoring. The combination of environmental specimen banking with metagenomic sequencing offers a powerful framework for uncovering hidden biodiversity trends in soil ecosystems and identifying the drivers of ecological reorganisation under global change.

## Background

Soil is among the most biodiverse habitats on Earth [1], supporting intricate communities of bacteria, fungi and invertebrates that underpin essential ecosystem functions such as nutrient cycling, carbon sequestration, and primary productivity [2,3]. Soil communities are key to terrestrial ecosystem health, yet their biodiversity patterns, especially temporal dynamics, are not well understood [4]. Long-term empirical data on soil biodiversity remain scarce [5], leaving crucial questions about the stability or change of belowground communities in the face of environmental pressures unresolved.

Large-scale biodiversity analyses have shown that local species richness (α-diversity) in many ecosystems remains relatively stable over time despite environmental change [6–8]. This stability is often attributed to species turnover: extirpations balanced by colonizations, rather than to stasis in community composition [9]. Consequently, changes in community composition (β-diversity) may occur even without shifts in richness. This decoupling has driven increasing attention to compositional homogenisation, i.e., the process by which communities become more similar across time or space [10–12]. Homogenisation may arise from environmental filtering, habitat simplification, or the spread of generalist species, and is considered a hallmark of directional ecological change [13,14].

To assess such compositional changes more precisely, abundance-based β-diversity can be decomposed into two components: balanced variation in abundance, which represents the reciprocal replacement of species (analogous to species turnover), and abundance gradients, which reflect uniform changes in abundance across all species (analogous to species nestedness where some communities are subsets of others) [11,15]. Empirical studies suggest that most temporal β-diversity is driven by balanced variation [7], but this may differ across habitats and taxonomic groups. In soil systems, such detailed analyses are still rare, especially across broad taxonomic scales encompassing bacteria, fungi, and metazoans.

A major barrier to understanding soil biodiversity change over time has been the lack of standardized, taxonomically broad time series. Soil communities are notoriously difficult to monitor due to their high richness, fine-scale spatial heterogeneity, and poor taxonomic resolution in traditional surveys [16,17]. However, environmental specimen banks provide an emerging solution [18]. The German Environmental Specimen Bank (German ESB) is a national long-term monitoring initiative in support of environmental management and policy [19] and has archived soil samples collected with uniform methods since 2002, offering a unique infrastructure for retrospective analyses [19]. Its cryogenically stored samples, coupled with modern metagenomic sequencing, now allow unprecedented insight into long-term biodiversity trends under natural environmental variability.

Here, we utilize 20 years of archived soil samples from 11 locations across Germany collected by the German ESBto examine temporal and spatial patterns of soil biodiversity across bacteria, fungi, and metazoans. Our study is based on high-throughput shotgun metagenomic data and aims to assess both α- and β-diversity using dissimilarity decomposition. Specifically, we hypothesize that (1) α-diversity has remained stable over time, reflecting resilience of local soil richness, and (2) β-diversity has shifted through compositional homogenisation, indicating directional change in community structure. By partitioning Bray–Curtis dissimilarity into balanced variation and abundance gradients, we explore the processes driving these patterns and examine whether temporal changes differ across taxonomic groups. Our study provides a rare empirical foundation for understanding how entire soil communities respond to environmental change, and highlights the value of environmental specimen banks for long-term ecological monitoring.

## Methods

### Study area

The sampling sites include 11 locations in nine sampling regions of different ecosystem types (conurbation, agrarian, forestry and near-natural) in Germany (Table 1). In addition to the primarily occurring forest vegetation, the study areas also consist of arable land in Bornhöved and lawn areas in Leipzig and Staden. The predominant soil type is brown soil, with exceptions being a rendzina in Berchtesgaden, a pseudovergleyed vega in Leipzig, a vega in Staden, and a pseudogley in Warndt. The average precipitation ranges from 4.7 mm (Leipzig) to 17.2 mm (Berchtesgaden) per month, and the average annual temperature ranges from 5.0 °C (Berchtesgaden) to 10.9 °C (Staden) (Table 1). Furthermore, the altitude varies from 35 m (Bornhöved) to 1,140 m (Berchtesgaden) above sea level. The dataset includes areas with various land uses. None of the soils from the sampling sites have been amended with any material from other locations, including sewage sludge. More detailed information on the individual sites can be found on the UPB website (https://www.umweltprobenbank.de/en) and in Table 1 and in Supplementary Figures S1-S11.

**Table 1.** Overview table of sampling sites.

| Study area | Code | Type of use | Source rock | Soil type |
| --- | --- | --- | --- | --- |
| Bayerischer Wald | BW | Forest | Granite, gneiss | Brown earth |
| Berchtesgaden | Bt | High mountain forest | Dolomite | Rendzina |
| Bornhöved | Bo | Arable land | Glaciofluvial deposits & alluvial deposits, sand alluvial deposits | Brown earths or colluvisol brown earths |
| Dübener Heide | Du | Forest | Ground moraine material | Strongly podsol brown earth |
| Harz | Ha | Forest | Granite | Brown earth |
| Leipzig | Le | Meadow | Alluvial sediments | Pseudogley-Vega |
| Pfälzer Wald | Pf | Forest | Red sandstone | Brown earth |
| Scheyern | Sy | Forest | Loess clay | Brown earth |
| Solling | So | Forest | Sandstone, loess clay | Brown earth |
| Staden | St | Meadow | Alluvial sediments | Vega |
| Warndt | Wa | Forest | Sandstone, flowing soils | Pseudogley from flowing soils on sandstone |

### Sampling

The samples were collected as part of the routine soil sampling of the German ESB [20]. As an integral part of the German pollution monitoring scheme, the German ESB sampled various environmental samples all over the country for multiple decades. The soil samples were collected using sterile equipment and according to highly standardised protocols [20]. Samples were frozen on liquid nitrogen immediately after collection, providing an unprecedented preservation of the sample associated biological community. The ESB’s soil sampling procedure has been standardized since 2002 and it is carried out every four years before leaf fall in autumn. Furthermore, the sampling areas were selected based on their representative reference soil type, a soil characterization of the area, and their proximity to terrestrial sampling sites.

Samples were taken from a 2,500 m^2^ sampling area. In forested sites, areas outside the canopy were avoided. Depending on soil texture and ecosystem type, litter layer (organic layer in ESB nomenclature), root network (the top soil layer from urban grasslands, including grass roots), or topsoil (A horizon) were sampled using a frame, spatulas and tweezers. The individual samples were then mixed horizontally outside the sampling area, weighed, and sieved to 2 mm. Excess material was either returned to the boreholes or deposited outside the sampling area. During sampling, a minimum weight of 5 kg was deep-frozen in stainless steel containers using liquid nitrogen. If necessary, the soil material was crushed using a jaw crusher, then mixed in a mixer, divided into sub-samples and stored as 50 g individual archive samples above liquid nitrogen for long-term storage in the ESB. The ESB provides cryogenic conditions in the archive with temperatures below approx. -130 °C (approx. < 140 °K), which is below the glass transition temperature of water. Furthermore, samples are stored above liquid nitrogen in an inert-gas atmosphere preventing oxidation processes. This ensures that biological and chemical processes in the samples are reduced to a minimum.

### Laboratory work & sequencing

Individual archive samples weighed under a clean bench. DNA was extracted with the ZymoBIOMICS™ DNA/RNA Miniprep Kit (Zymo Research, Freiburg, Germany). Negative controls were included during extraction to check for contamination. The manufacturer’s protocol was followed with the following modifications: before homogenization, an additional mechanical grinding step (5 min) was performed using three 3 mm beads at 30 Hz on the Retsch mill. This was followed by further beating (15 min) in BashBead Lysis Tube. DNA concentrations were quantified using the QuantiFluor® ONE dsDNA System (Promega, Madison, USA). Fragment lengths were assessed with a TapeStation 2200 (Agilent Technologies, Waldbronn, Germany). The concentrations of individual samples were normalised, and the extracts were fragmented using the Covaris M220 Focused-ultrasonicator for 85 sec (Covaris, Massachusetts, USA). Libraries were prepared and purified according to the BEST protocol [21]. Fragment lengths were checked again with an agarose gel electrophoresis before pooling the samples into two pools with separated replicates. Sequencing was performed on an Illumina NovaSeq 6000 PE150 platform at Novogene (Cambridge, UK), with a sequencing depth of 2 GB per sample.

### Bioinformatics processing

Sequences were automatically trimmed and quality checked with Autotrim v0.6.1 and FastQC. Autotrim depends on Trimmomatic [22], and MultiQC [23] to trim, screen for overrepresented k-mers and create summary statistics, respectively. Taxonomic classification was performed sequentially using Kraken2 v2.1.2 [24] against several reference databases with a confidence interval of 0.95. For building the reference databases, representative or reference RefSeq assemblies were downloaded from NCBI (downloaded on January 13, 2023) for the following RefSeq categories: human, bacteria, and fungi and plants combined. For invertebrates, the newly created MetaInvert database was used, which contains genomes of 232 Central European invertebrate soil species [25]. Each reference dataset was used to build a Kraken2 database with the default k-mer size (k = 35). For the sequential analysis, the unclassified sequences of the previous classification were conducted against the following database with the order mentioned before and the MetaInvert database used as the last reference database. Results of the sequential runs were merged in a custom script. Raw sequencing data is available in NCBI under accession number **XXX**.

### Data analysis

The data were analysed with R v4.4.3. If read counts of a taxon in a sample exceeded at least five times the maximum read count of that taxon in any negative control, we subtracted the control reads from the sample reads. Otherwise, the taxon was entirely removed from that sample. The dataset was grouped into three groups for analysis: bacteria, fungi, and metazoa. Unclassified reads were removed. We considered a sample for analysis only of the taxonomic richness of any of three groups was higher than 10. All analyses were performed with taxa at the best possible taxonomic identification.

Analysis of biodiversity and community composition trends were performed with a custom script (PRSB_analysis_script.R) with vegan v2.6-6.1 [26] and mvabund v4.2.1 [27]. We calculated temporal Bray-Curtis dissimilarities among temporal samples of each of the 11 sites to evaluate temporal homogenisation in community composition. In addition, we decomposed Bray-Curtis dissimilarities into a component accounting for balanced variation in abundance and abundance gradients [11]. Balanced variation in abundance represents the replacement of individuals of species with individuals of different species. Abundance gradients represent the overall loss or gain of individuals at a site compared to each other across all species. Balanced variation in abundance and abundance gradients are the conceptual analogues of turnover and nestedness components of incidence-based β-diversity metrics [15]. We evaluated temporal and spatial trends in α- and β-diversity with analyses of variance of linear models. Linear models of α- and β-diversity were fit with total reads, sampling year and sampling locality as predictors. We included the total number of reads obtained per sample into models to account for eventual differences caused by uneven sequencing depth. This resulted in a conceptual model formula of diversity ∼ reads + year + locality.

We evaluated temporal and spatial patterns of community composition with multivariate generalized linear models (GLMs) for abundance data [27]. Multivariate GLMs fit separate regressions on each species, without a need for normalizing count data [28]. They also do not confound location and dispersion effects, a typical issue of distance-based compositional comparisons such as PERMANOVA [29]. These properties make multivariate abundance models highly suitable for analysing community composition recorded with high throughput sequenced DNA reads, a data type known for non-normal distribution and overdispersion [30]. We assumed that read counts were generated with a negative binomial distribution which is considered reasonable for metagenomic data [31]. Multivariate abundance models were fit with total reads, sampling year and locality or land use as predictors (diversity ∼ reads + year + locality or diversity ∼ reads + year + land use). We tested individual predictor effects with a likelihood-ratio-based analysis of variance for multivariate GLMs, with 999 bootstrap iterations, Monte-Carlo resampling. We summed AIC values of individual species-specific models to compare models fit with locality or land use as spatial predictors. We selected core taxa for community composition analyses of each of the three groups [32]: we modelled the log-normalized average read abundance of taxa against their incidence (the number of samples in which a particular taxon is present) with generalized additive models (GAM) with the mgcv package v1.9-1 [33]. GAMs are highly suitable for exploring incidence-abundance patterns as they do not impose a strict parametric relationship between variables [24]. We included bacterial taxa if they were present in more than five, fungal taxa in more than eight and metazoan taxa in more than four samples based on the GAM-identified patterns.

## Results

DNA extracts were obtained from 55 bulk soil samples, covering 11 locations and 20 years, at 4-year intervals. A total of 1.4 billion reads were sequenced, of which 99.55 % could not be assigned to any taxa. Of 0.6 million post-processed reads, approximately 51% were assigned to Viridiplantae, 34% to Bacteria, 13% to Metazoa, and 3% to Fungi. Total α-diversity of bacteria, fungi and metazoa ranged between 50 (Harz in 2002) and 1449 taxa (Bornhöved in 2006) during 20 years of ESB sampling, with no statistically significant trends over time for the overall dataset (Fig. 1, Table 2). Taxonomic richness was highest at arable sites for all groups (Fig. 2).

**Table 2.**
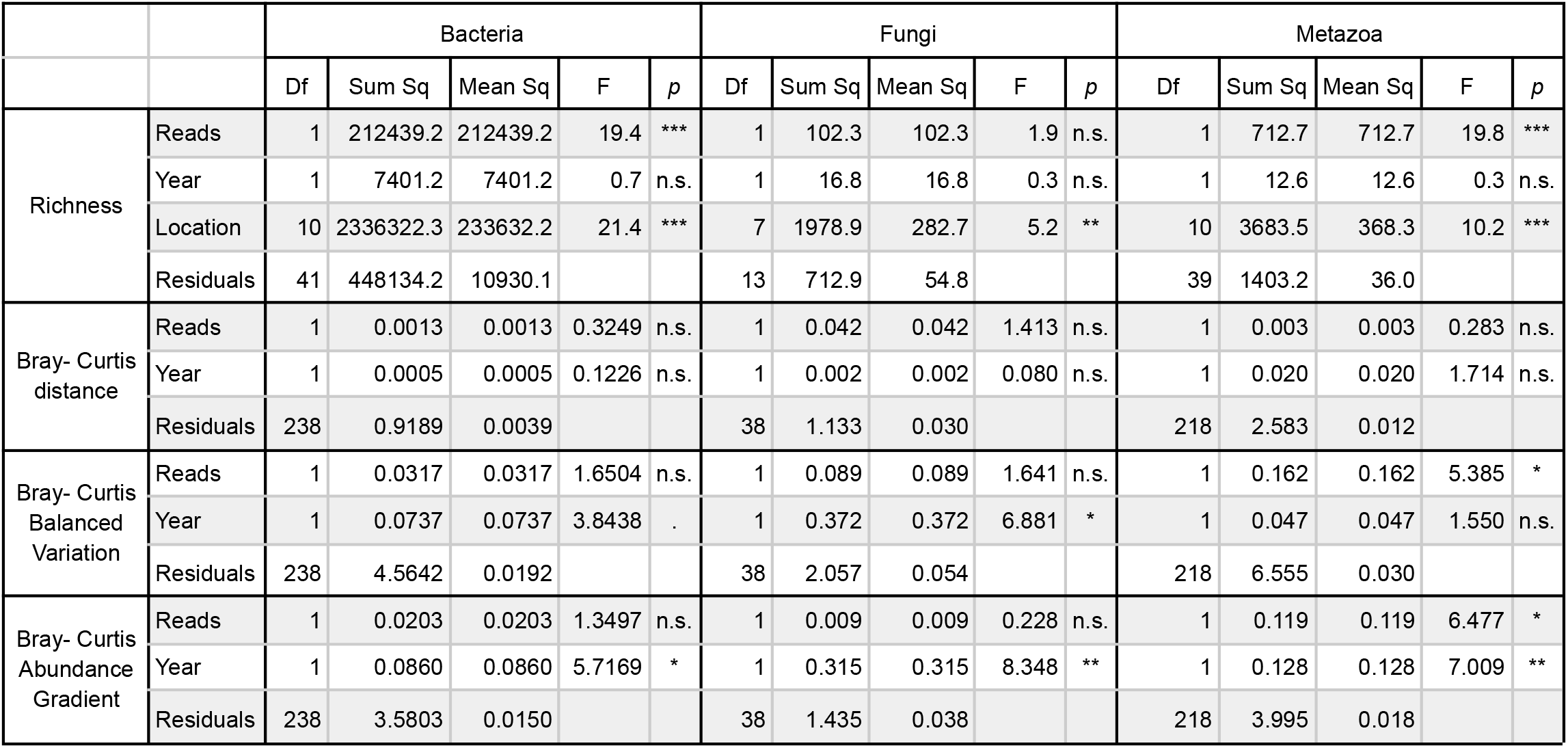
Analysis of variance tables of spatial and temporal patterns of α- and β-diversity based on linear models. Statistical significance: *** *p* ⩽ 0.001, ** *p* ⩽ 0.01, * *p* ⩽ 0.05,. *p* ⩽ 0.1, n.s. not significant.

**Fig. 1:**
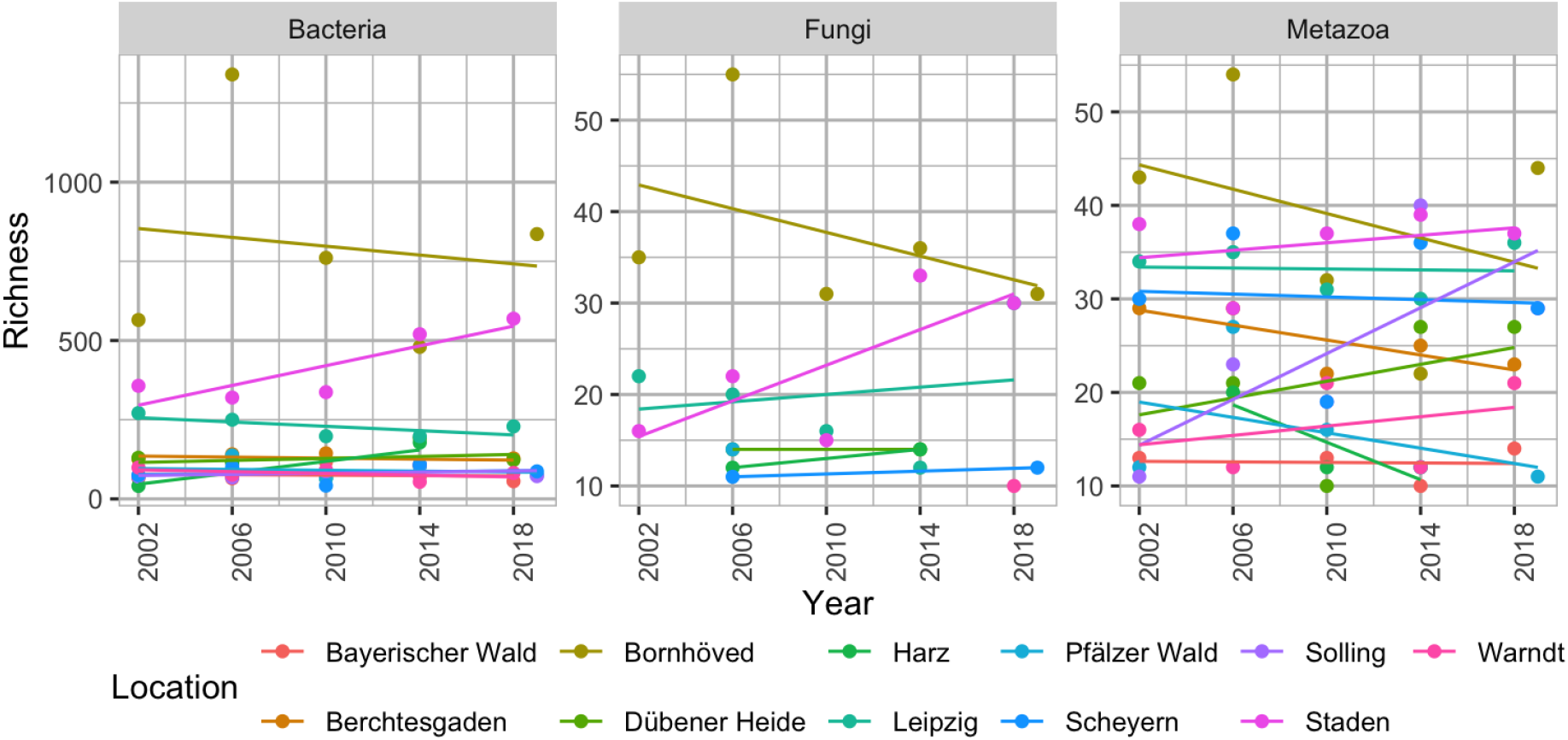
Temporal trends of α-diversity for the different organism groups and locations with lines representing linear trends. Dots show taxon numbers colored according to locations, grouped by year of sampling y. Lines mark linear trends.

**Fig. 2:**
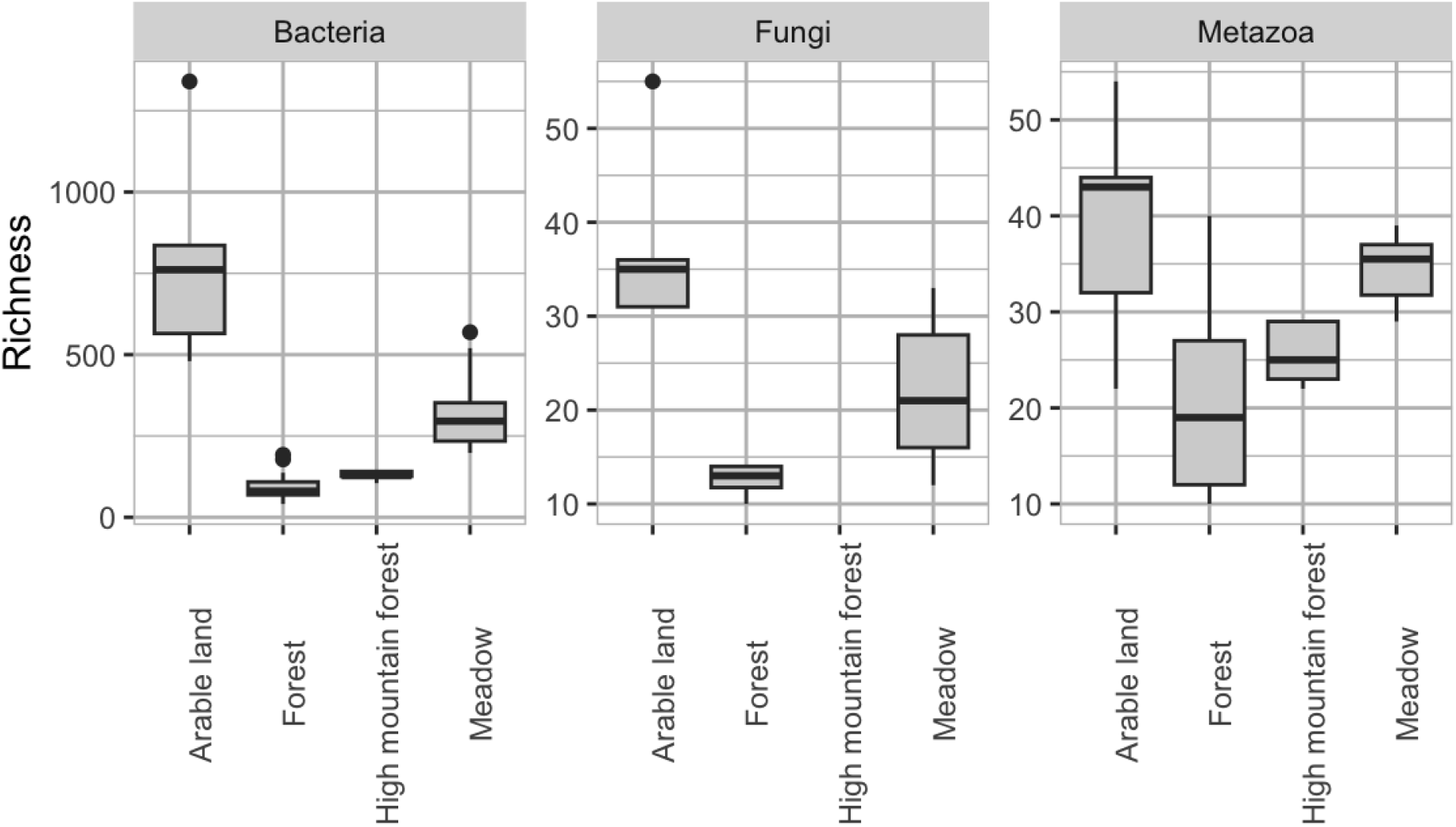
Taxonomic richness of organism groups at different land uses. Boxplots show median, lower and upper hinges (first and third quartiles) and whiskers (extending from the hinge to the smallest or largest value no further than 1.5 times the distance between the first and third quartiles). Data beyond are considered outliers and are plotted individually.

Trends of richness and Bray-Curtis community dissimilarity were stable through time for all groups (Fig. 3, Table 2). The balanced component of temporal turnover marginally significantly decreased for bacteria, and statistically significantly decreased for fungi. The gradient component of temporal turnover statistically significantly increased for bacteria and fungi, and decreased for metazoa (Fig. 3, Table 2).

**Fig. 3:**
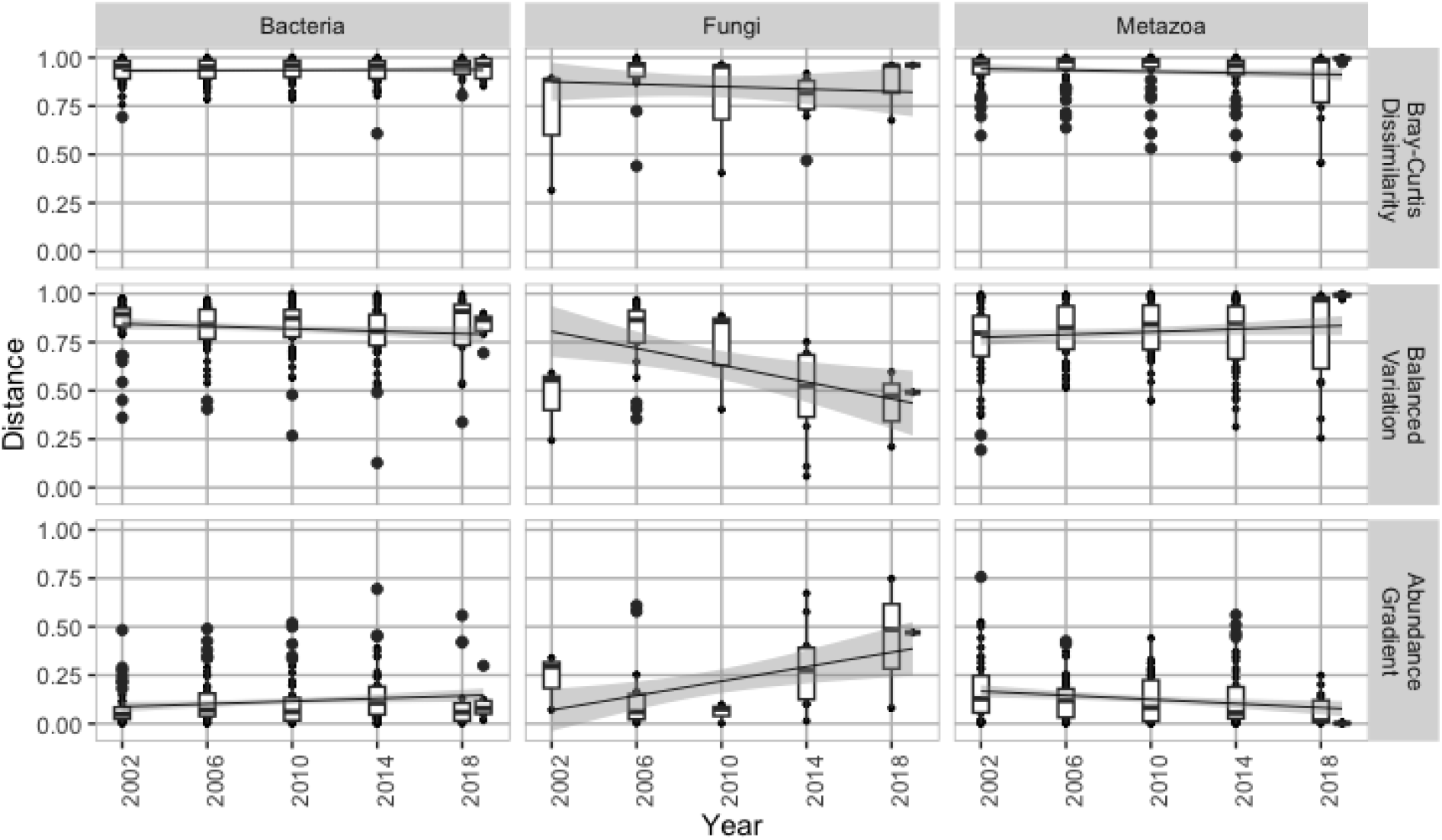
Temporal trends in Bray-Curtis dissimilarity and its balanced variation (analogues of turnover) and gradient components (analogues of nestedness). Boxplots show median, lower and upper hinges (first and third quartiles) and whiskers (extending from the hinge to the smallest or largest value no further than 1.5 times the distance between the first and third quartiles). Data beyond are considered outliers and are plotted individually.

There were no statistically significant temporal differences in community composition (Fig. 4, Table 3). Community composition was statistically significantly defined by locality or land use. Land use as spatial predictor resulted in a better model fit for bacteria (∑AIC_locality_ = 57,499, ∑AIC_landuse_ = 56,838) and fungi (∑AIC_locality_ = 2,758, ∑AIC_landuse_ = 2,749), while model with with locality as spatial predictor was better for metazoa (∑AIC_locality_ = 11,507, ∑AIC_landuse_ = 11,725).

**Table 3.** Analysis of variance tables of multivariate abundance GLMs of community composition.

| Spatial predictor |  | Bacteria |  |  |  | Fungi |  |  |  | Metazoa |  |  |  |
| --- | --- | --- | --- | --- | --- | --- | --- | --- | --- | --- | --- | --- | --- |
|  |  | Res.Df | Df.diff | Dev | *p* | Df.diff | Df.diff | Dev | *p* | Res.Df | Df.diff | Dev | *p* |
| Location | Intercept | 53 |  |  |  | 47 |  |  |  | 53 |  |  |  |
|  | Reads | 52 | 1 | 807 | *** | 46 | 1 | 26.1 | 0.073 | 52 | 1 | 124.1 | ** |
|  | Year | 51 | 1 | 460 | n.s. | 45 | 1 | 34.4 | 0.009 | 51 | 1 | 83.0 | n.s. |
|  | Location | 41 | 10 | 18120 | *** | 35 | 10 | 681.5 | 0.001 | 41 | 10 | 2617.2 | *** |
| Land use | Intercept | 53 |  |  |  | 47 |  |  |  | 53 |  |  |  |
|  | Reads | 52 | 1 | 807 | *** | 46 | 1 | 26.1 | 0.053 | 52 | 1 | 124.1 | ** |
|  | Year | 51 | 1 | 460 | n.s. | 45 | 1 | 34.4 | 0.008 | 51 | 1 | 83.0 | n.s. |
|  | Land use | 48 | 3 | 11166 | *** | 42 | 3 | 438.5 | 0.001 | 48 | 3 | 1223.6 | *** |

**Fig. 4:**
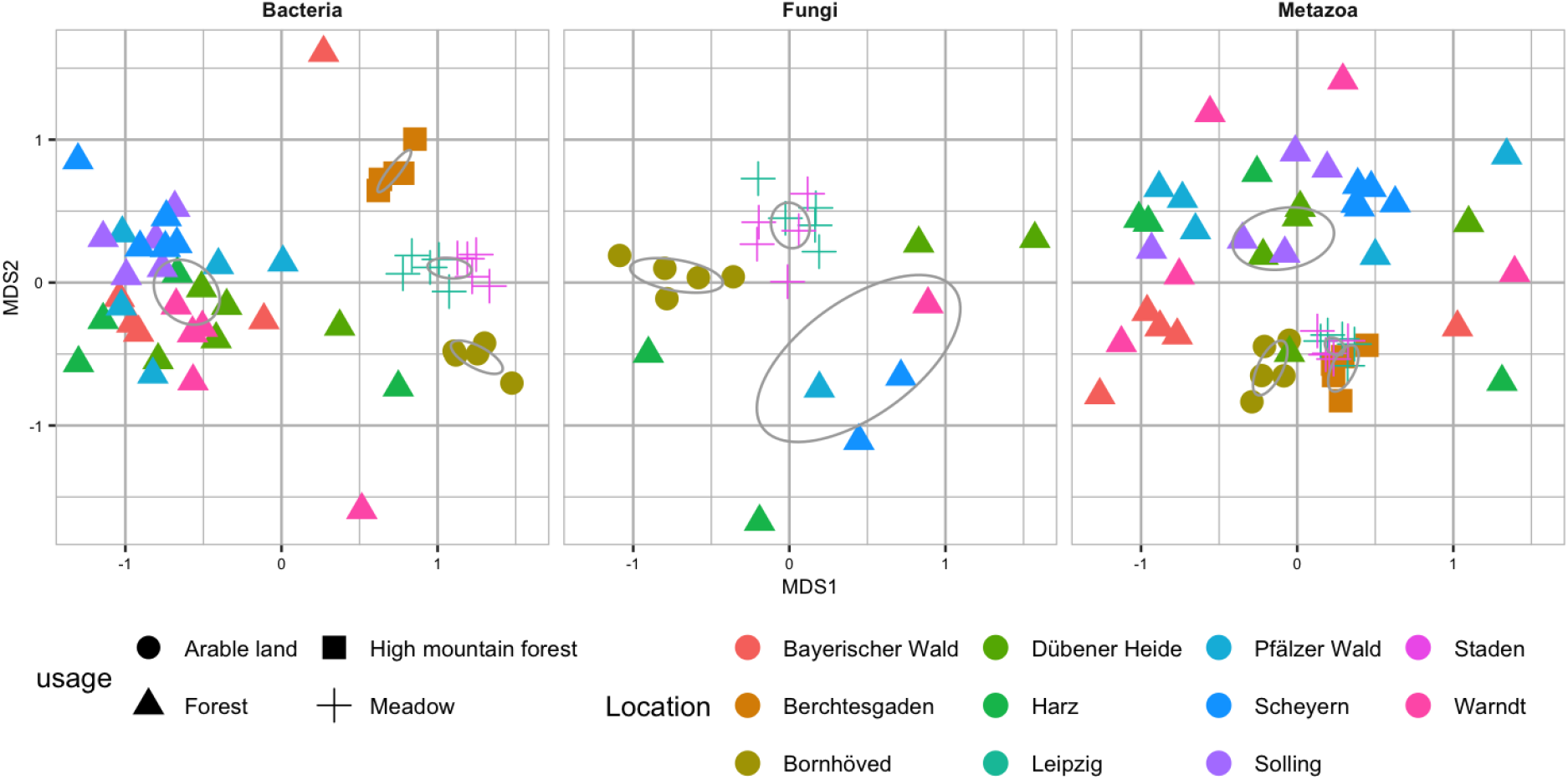
Non-metric multidimensional scaling ordination of bacterial (stress: 0.164), fungal (stress: 0.17) and metazoan (stress: 0.259) community composition based on Bray-Curtis dissimilarities. Grey ellipses denote the 95% confidence interval of land use group centroids.

## Discussion

### No temporal α-diversity trends, but strong spatial structuring

Our long-term data show no significant change in soil α-diversity for bacteria, fungi, or metazoa over the 20-year study period (Fig. 1). This finding aligns with broader patterns observed in other ecosystems where local species richness often remains stable through time [6,7]. In those studies, the lack of a net diversity trend is attributed to species replacements balancing out species losses, rather than an absence of community change. Similarly, in our soil communities, it appears that local extinctions may have been offset by colonization or resurgence of other taxa, resulting in relatively constant richness. In contrast to the lack of temporal trends, we observed pronounced spatial differences in diversity. Some locations consistently harbored far more taxa than others (ranging from ∼50 to nearly 1,500 observed taxa in extreme cases), indicating that site-specific factors strongly influence how much biodiversity the soil can support. Such spatial heterogeneity in soil biodiversity is well documented and often reflects underlying differences in soil properties, climate, and habitat type across locations [2]. The observed spatial patterns support that long-term environmental conditions at the origin of a sample are major determinants of species richness, often overpowering shorter-term differences [34].

Our results revealed pronounced differences in soil taxon richness across land-use types, with the strongest contrasts observed in bacteria. Bacterial richness peaked in arable land, while forests and high mountain forests consistently supported lower bacterial diversity (Fig. 2). This pattern mirrors recent observations across Europe [35,36], where intensively managed agricultural soils often harbor more bacterial taxa than less disturbed habitats. The increased richness at arable sites might be a result of increased resource (nutrient) availability which is contributing to larger population sizes across taxa, and consequently to higher richness [37]. Both fertilisation and tillage increase the accessibility of nutrients to organisms, potentially causing the observed pattern [36].

### Stable total β-diversity conceals taxon-specific compositional shifts

Bray-Curtis dissimilarity was consistently high among the localities, and it showed no temporal trends (Fig. 3, Table 2). However, when separated after location, the balanced variation and abundance gradient components of Bray-Curtis dissimilarity [11] revealed dynamic temporal reorganization. The balanced variation component dominated compositional dissimilarity in comparison to the abundance gradient component. This is similar to temporal β-diversity trends recorded for aquatic and terrestrial taxa, where temporal turnover dominated over nestedness in 97% of the analysed time series [7].

Balanced variation and abundance gradients showed divergent patterns among bacteria, fungi, and metazoans. Over the 20-year period, bacterial and fungal communities exhibited increasing abundance gradients, coupled with relatively stable or slightly declining balanced variation (Fig. 3; Table 2). These patterns are consistent with compositional homogenization, where communities become increasingly dissimilar due to general shifts in taxon abundances, rather than species turnover. In contrast, metazoan communities showed no signal of homogenization. Instead, we observed declining abundance gradients and stable balanced variation, suggesting greater temporal stability in composition. This highlights a fundamental difference in the temporal dynamics and ecological sensitivities of microbial *versus* faunal soil communities. For bacteria and fungi, the observed changes may reflect growing dominance of tolerant or opportunistic taxa, potentially favored by environmental filtering, such as pollution, fertilization, climate trends, or vegetation shifts [13,14], and increased availability of nutrients and energy [37]. Such differentiation is a signal of directional environmental change, particularly where functional redundancy permits reordering rather than extinction [12]. In contrast, the decline in abundance gradients by metazoans with no significant changes in balanced variation indicates increased evenness across time and more stable abundance distributions. In contrast, metazoan communities showed no signs of homogenization or differentiation. This may reflect ecological inertia resulting from the slower life histories of metazoans compared to microorganisms. Other relevant processes include dispersal limitation [38], and stochastic community dynamics predicted by neutral metacommunity models [39].

While overall Bray–Curtis dissimilarities among sites remained temporally stable, this apparent equilibrium masked substantial compositional restructuring within communities within sites. This reflects a common challenge in biodiversity monitoring: aggregated metrics often obscure dynamic ecological processes [9,10,40]. In our case, the temporal decomposition of Bray-Curtis dissimilarity into balanced variation and abundance gradients revealed clear taxon-specific signatures of change. These two components reflect fundamentally different ecological processes [11,41], and working with these components of dissimilarity allowed us to detect subtle but ecologically meaningful reorganization in soil communities. This underlines the importance of multifaceted analyses to reveal divergent biodiversity trajectories that would be missed by summary metrics.

### Land Use and Locality as Drivers of Soil Community Composition

Both the broad land-use category of a site and its specific locality (geographic location and context) shaped soil community composition, but their relative importance differed among organism groups. For bacteria and fungi, land use explained more variance in community composition than locality. This indicates that the type of habitat and management regime (e.g. forest, meadow, arable) was a strong determinant of microbial communities. This makes sense since land use influences microbes through key environmental filters [2] such as soil nutrients [42], organic matter inputs [43], disturbance frequency [44], and vegetation type [45]. Indeed, land-use has been recognized as a consistent driver of soil microbial community assembly at regional scales [46], often overriding pure spatial distance. Our findings support this: two forest sites in different parts of Germany may foster more similar bacterial communities than a meadow and a forest located at shorter distances (Fig. 4), because similar soil conditions select for certain microbes. This pattern aligns with the classic hypothesis of Baas Becking from 1934 that “everything is everywhere, but the environment selects”: many microbial taxa can disperse well, and it is the local habitat, strongly tied to land use, which determines community composition [47]. In contrast, the composition of soil metazoan communities was better explained by locality than by broad land use type. Soil animals often have limited dispersal ranges and specific microhabitat requirements, which, adding to the effects of environmental filters [3], can lead to more pronounced spatial turnover even within the same land use across short spatial distances [16]. While environmental factors strongly influence soil metazoan communities [3,48], our results also support the importance of historic biogeography and dispersal limitation on compositional differences. The differences highlight that land management for soil biodiversity may need to be taxon-specific: while the effects of habitat alterations may be more predictable for microbes and fungi, managing soil animal diversity might require more locally tailored strategies.

### Perspectives

Metagenomics is regularly used for assessing the composition and function of microbial communities, but so far it is rarely used for metazoans [49]. A chief challenge is the taxonomic assignment of sequences: in our dataset a staggering 99.5% of raw reads could not be classified to any known organism. This reflects the incomplete state of reference databases not only for soil invertebrates [25], but also for soil bacteria [50]. Consequently, our diversity estimates rely on the minority of reads, assigned to well-studied groups. It is likely that the true diversity is higher than we report. Community changes, especially among poorly characterized microbes or microfauna, might go undetected due to this “dark matter” of unclassified reads [50]. Another limitation is that metagenomic data do not distinguish between live organisms and residual environmental DNA [51]. Soil can retain DNA of organisms long after their death or dormancy, which means detections might include historical remnants. This could actually inflate stability in a long-term study, as past community members’ DNA lingers. Simultaneous use of eDNA and eRNA might help to mitigate legacy effects due to short eRNA degradation times [52].

Nevertheless, high-throughput metagenomics offers several benefits for biodiversity monitoring. First, it provides a comprehensive view of life in soil, from microbes to multicellular organisms compared to other approaches such as taxon-specific metabarcoding. Metagenomics evaluates all organisms with a single method, without needing multiple primers, each with its own amplification biases [53]. Second, the lack of taxon-specific marker gene amplification offers a less biased way of quantifying read abundances [49]. This allows for a semi-quantitative insight into community structure, beyond species presence [54]. Finally, the lack of a marker-gene-specific PCR makes metagenomics more replicable and standardisable in comparison to metabarcoding, where primer choice is one of the key factors which influence recorded community composition [55].

Working with decades-old samples poses logistical challenges: DNA may be degraded if not stored properly [56], although soil eDNA preservation might be remarkably stable [57]. We were fortunate to have well-preserved samples and applied uniform methods. Tapping into ESB resources opens up high-quality, well-documented sample collections that allow easily adding a retrospective component to biomonitoring projects. This underlines the importance of environmental specimen banks and sample collections in long-term biodiversity research. Spatial monitoring programs should consider retaining samples for future investigation, while existing ESBs should consider expanding their sampling and archiving strategy to include soil samples - if they are missing. Recent developments in soil policy (e.g. the EU Soil Strategy [58]and a proposed Directive for Soil Resilience and Monitoring [59] trigger national awareness for better protection of soil, preventing and remediating contaminated soils and implementing soil biodiversity monitoring. Discussions have started on the design of regulatory monitoring programs and baseline studies on the EU and national levels, and these studies can provide systematic overviews on spatial and temporal soil biodiversity [60]. Robust biodiversity data for soil is often lacking, especially for systematic trend comparisons. Retrospective monitoring data from existing soil collections and ESBs can at least partially fill the missing historical context when temporal trends are linked to a denser network of spatial data from new monitoring programs. In addition, retrospective studies can inspire emerging monitoring programs by testing novel methods, e.g. DNA-based biodiversity monitoring. Thus, the data from this study can provide important insights into the robustness and plausibility of the data for future trend observations as well as first information on the temporal variability of the observed taxonomic groups.

Our study revealed temporal trends in regional soil biodiversity, but it did not address the actual drivers of community change. Future studies should investigate and quantify the effects of specific factors, such as pollution, land use change and global warming. Linking drivers to biodiversity trends will need a more extensive spatial sampling scheme. This might be possible by joining efforts with other national ESB initiatives [61,62] which are collecting and preserving samples from other regions. A combination of time series data with spatially extensive time-for-space sampling [63] might be promising for disentangling collinear drivers of soil biodiversity change.

## Conclusions

In conclusion, our study demonstrates that, while aggregated diversity indices may suggest stability in soil biodiversity over time, they obscure significant underlying compositional restructuring, particularly among microbial taxa. Using a uniquely standardized, long-term environmental sample archive we uncovered consistent patterns of directional change in soil communities, likely driven by shifts in taxon abundances rather than species turnover. These findings underscore the need to move beyond aggregated richness metrics and adopt nuanced, taxonomically inclusive approaches to biodiversity monitoring. The differing trends of bacteria, fungi, and metazoa also highlight that soil biodiversity does not respond uniformly to environmental change. As land-use intensification continues to alter soil environments, understanding taxon-specific dynamics will be crucial for targeted conservation and land management strategies. Future efforts should aim to integrate environmental metadata, functional trait information, and broader spatial sampling to better resolve the drivers behind these hidden biodiversity shifts. Long-term metagenomic monitoring, especially when coupled with specimen banking, holds immense promise for capturing the complex and often cryptic dynamics of soil ecosystems under global change.

## Acknowledgements

We thank Karl-Heinz Weinfurtner, Fraunhofer Institute for Molecular Biology and Applied Ecology (IME) for collecting the soil samples and Dr. Bernd Göckener and ESB team at Fraunhofer IME for long-term archiving the soil samples. We thank Róbert Veres for generating and curating the reference genome databases used for the taxonomic assignment of the metagenomic data. The work was funded by the TrendDNA project of the German Federal Environment Agency (Umweltbundesamt). Additional funding to MB was provided by the DFG Priority Program 1374 “Biodiversity Exploratories” (BE-Spring BA 4843/4-1, MA 7144/2-1, SCHN 1620/1-1), and the Landes-Offensive zur Entwicklung Wissenschaftlich-ökonomischer Exzellenz (LOEWE) Program of the Hessian Ministry of Higher Education, Research, Science and the Arts through the LOEWE/1/10/519/03/03.001(0014)/52 project (LOEWE Centre for Translational Biodiversity Genomics - LOEWE-TBG).

## Use of Artificial Intelligence (AI) and AI-assisted technologies

AI-assisted tools (NotebookLM, ChatGPT, Perplexity, DeepL Write) were used to improve the readability of the text, and as search engines. These tools were used solely to improve the authors’ original research content. All scientific insights, data analysis, interpretations, and conclusions were conceived and performed by the authors. The authors take full responsibility for the accuracy, originality and integrity of all content.

## Data availability statement

xxx

## Author contributions

Conceptualisation: JP, HK, MB; Data curation: JR; Formal analyses: JP, JR, MB; Funding acquisition: MB; Investigation: JP, JR, KT, HK, MB; Methodology: JP, JR, DB, LB, LS, HK, MB; Project administration: HK, MB; Resources: LS, HK, MB; Supervision: KT, HK; Validation: KT, MB; Visualisation: JP, MB; Writing – original draft: JP, KT, MB; Writing – review & editing: all authors.

