## Supplementary Figs S1-S11 for "Two decades of compositional restructuring of soil biodiversity in Germany despite stable α- and β-diversity indices"

### Supplementary Materials

#### German Environmental Specimen Bank: land use types and spoil profiles

Figs. S1 - S11: Soil profiles from the 11 sampling sites

**Arable land**

Bornhöved: The Bornhöved Lake District is in the main water divide between the North- and Baltic Sea. The land is mainly used for agriculture. The sampling site Ruhwinkel-Ost is located on arable land in the catchment area of Lake Belau.

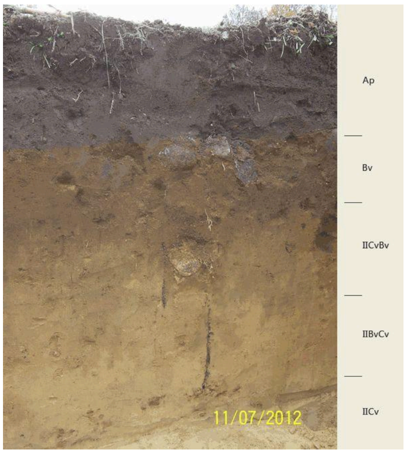


Fig. S1: Soil profile in Bornhöved

#### Forest

Bayerischer Wald (Großpalmberg): The Upper Bavarian Tertiary Uplands are a part of the Southern German Molasse Basin. There are only a few large contiguous forest areas. The majority of the forest is small scaled and highly parcelled. Outside the forest, the land is used primarily for agriculture. The sampling site Großpalmberg is located in a mixed forest.

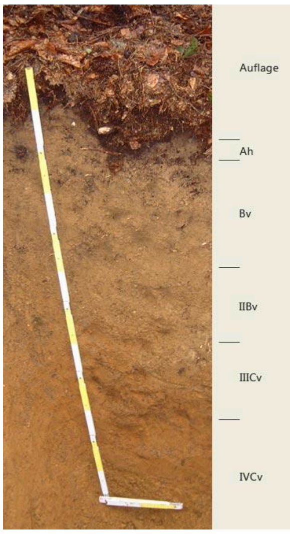


Fig. S2: Soil profile at Bayerischer Wald (Großpalmberg)

Solling: Second highest and largest low mountain range in Northern Germany. The sampling site Friedrichshäuser Bruch is located in the forest district Sievershausen. The soil sampling site is situated in a spruce stand.

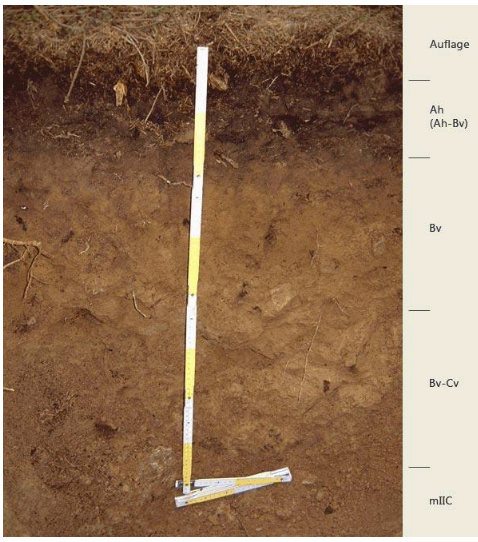


Fig. S3: Soil profile in Solling

Pfälzer Wald (Palatinate Forest): Germany's largest connected forest area in a range of low mountains. The soil sampling site Edersberg is situated in a beech stand.

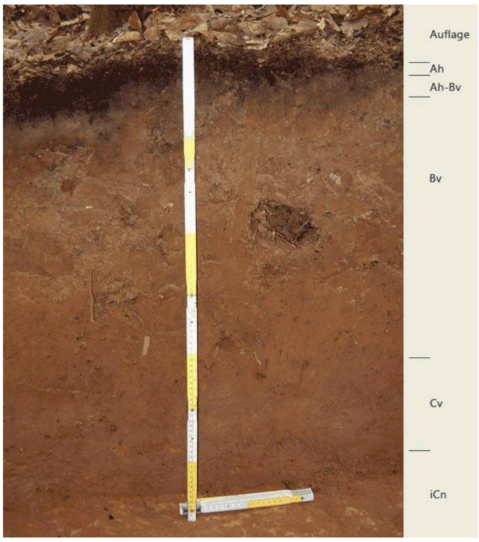


Fig. S4: Soil profile in the Pfälzer Wald

National Park Harz: Germany's largest forest national park. The sampling site Naturdenkmal Stempelsbuche is located in a spruce forest.

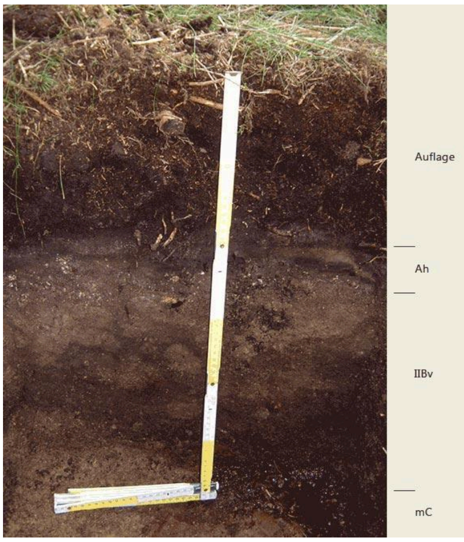


Fig. S5: Soil profile in the Harz

Scheyern (Bayerischer Wald): Germany's first national park with extensive forests. The dominant tree at this site is the beech.

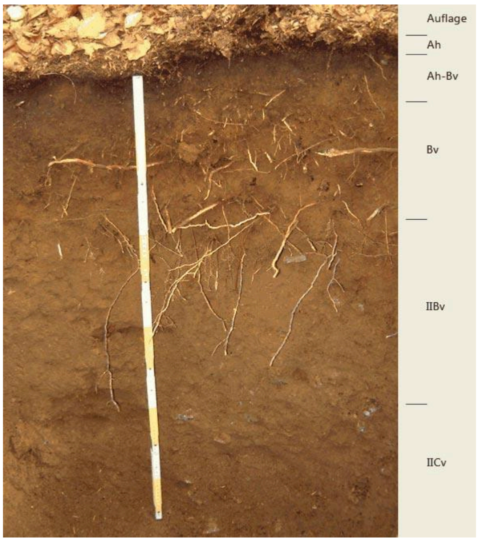


Fig. S6: Soil profile at Scheyern (Bayerischer Wald)

Dübener Heide: One of the largest contiguous forests in the region and an important recreational area of the Halle-Leipzig conurbation. The sampling site Revier Lutherstein is situated in a beech stand.

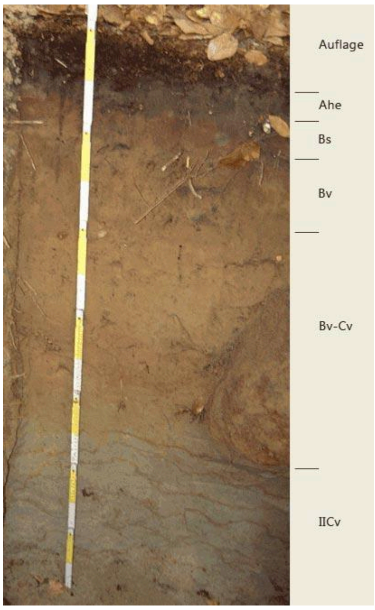


Fig. S7: Soil profile at the Dübener Heide

Warndt: Forest ecosystem between the industrial regions of the Saarland and Lorraine. The sampling site Warndt 2 is located in the forest district Warndtweiher. The soil sampling site is situated in a beech mixed forest.

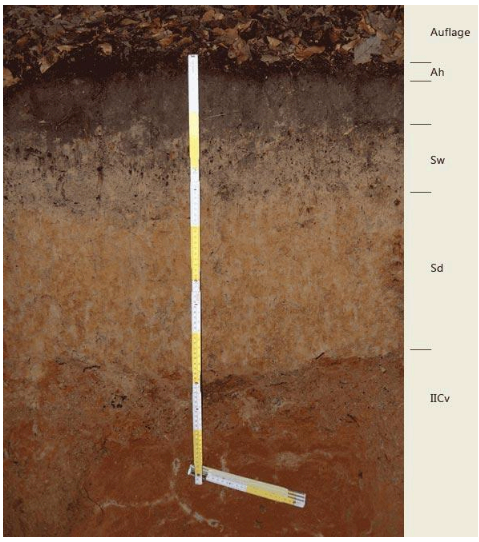


Fig. S8: Soil profile Warndt 2 (Site U2-B); analysed sample is the litter layer (“Auflage”)

##### High mountain forest

Berchtesgaden: The only high mountains national park in Germany and an area of the Limestone Alps with international relevance. The soil sampling site Wimbachtal is situated in a spruce stand.

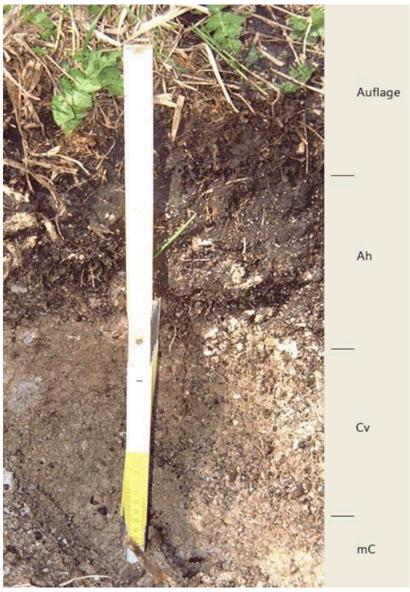


Fig. S9: Soil profile in Berchtesgaden

**Meadows**

Leipzig, Rosental: Grassy park area in the centre of Leipzig city.

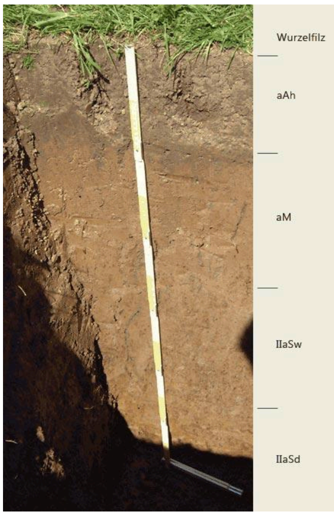


Fig. S10: Soil profile in Leipzig

Staden: The sampling site is located in a grassy floodplain between the Saar and the city of Saarbrücken and is used as a park area.

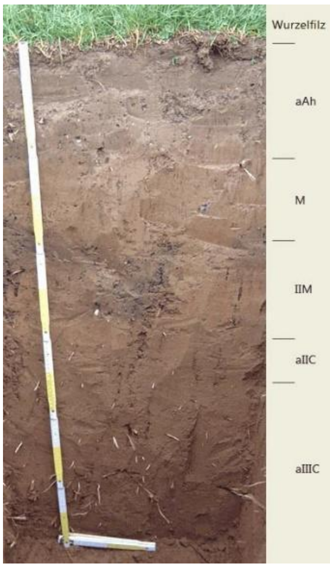


Fig. S11: Soil profile in Staden
